# A machine learning approach to infer DNase1L3 activity from plasma cell-free DNA fragmentomics

**DOI:** 10.64898/2026.03.16.712205

**Authors:** Jasper Linthorst, Erik A. Sistermans

## Abstract

DNase1L3 is an endonuclease that fragments DNA during apoptosis and digests DNA from microparticles in plasma, shaping key features of cell-free DNA (cfDNA) and maintaining extracellular DNA homeostasis, a process implicated in autoimmunity. The common missense variant p.Arg206Cys (R206C) affects cfDNA through a non-linear allele dosage effect, with limited effects in heterozygotes and strong effects in homozygotes. Therefore, commonly used models trained on fragmentomics from individuals with normal DNase1L3 activity perform poorly in R206C homozygotes.

To address this, we analyzed cfDNA sequencing data from 129,676 Non-Invasive Prenatal Tests and validated R206C genotypes in a selection of 169 matching plasma samples. Supervised and unsupervised learning were used to infer DNase1L3 activity from cfDNA fragmentation properties.

Our models accurately identify R206C homozygotes using as little as 10,000 cfDNA fragments, outperforming genotype imputation. However, unsupervised analysis reveals few samples that cluster with homozygotes but lack the corresponding genotype, suggesting that our method also identifies other or downstream effects of DNase1L3 impairment. Conversely, some R206C homozygotes initially lacked the aberrant fragmentome, but longitudinal follow-up across subsequent pregnancies shows that they develop aberrant fragmentomes over time.

These findings enable the identification of samples with impaired DNase1L3 activity directly from sequencing data, providing a practical approach to improve interpretation and robustness of cfDNA-based diagnostics across clinical applications.

## Introduction

Cell-free DNA (cfDNA) in plasma has become an essential biomarker in non-invasive clinical applications, including prenatal screening, cancer diagnostics, cardiology and organ transplant monitoring^1^. The intra- and intercellular cleavage of DNA by different nucleases gives rise to the so-called fragmentomic features of cfDNA, which include specific fragment sizes, nucleosome footprints, end motifs, and other nucleotide patterns. These molecular properties of cfDNA fragments are of great interest for many different applications of non-invasive diagnostic-en screening tests. Predominantly, but not exclusively, cancer diagnostics. However, the biology behind these fragmentomics signatures is not completely understood. Recent work has shown that the fragmentomics signatures of cancer largely resemble those of other diseases, such as venous thromboembolism and autoimmunity^2^. This suggests that these patterns may all stem from shared underlying biological processes.

DNase1L3, a member of the DNase I family of nucleases, plays a crucial role in the fragmentation of DNA during apoptosis and is essential for the efficient clearance of extracellular DNA in plasma. Although different nucleases are known to be involved in the generation of cfDNA, DNase1L3 was shown to account for a large fraction of the fragments in plasma^3,4^. Loss-of-function variants in DNase1L3 cause a rare monogenic form of systemic lupus erythematosus (SLE)^5^. The more common p.Arg206Cys missense variant (R206C) has been shown to increase the risk of a spectrum of other autoimmune diseases, including SLE, Rheumatoid Arthritis and Systemic Sclerosis^6^. R206C affects the folding^7^ and/or secretion^8^ of DNase1L3, which reduces the activity of the protein. Mechanistically, this is linked to the capacity of DNase1L3 to release microparticle-associated DNA^9^ and degrade neutrophil extracellular traps^10^ (NETs), a process which also plays an important role in shaping the tumor microenvironment^11^.

The genomic positions where nucleases such as DNase1L3 cleave is largely determined by DNA binding proteins, specifically nucleosomes. The nucleosomal organization is highly cell-type specific, and because most cfDNA is still bound to nucleosomes, the fragmentomic features of cfDNA can be used to determine a differential contribution of tissues or cell-types to the pool of cfDNA^12^. Typically, variability in these fragmentomic features is attributed to factors that drive this altered contribution, such as the turnover of placenta or tumor-derived cells. However, this wrongfully assumes consistent nuclease efficiency across individuals.

We previously conducted a large Genome Wide Association Study (GWAS) to explore the genetics of cfDNA and its fragmentomic characteristics. Our findings indicated that the previously mentioned R206C variant in DNase1L3, which has a minor allele frequency of around 7% in the European population, serves as a key genetic factor influencing the variability of cfDNA properties^13^. With a non-linear allele dosage effect the variant decreases the number of mononucleosome-sized fragments and causes a relative increase in multi-nucleosome sized fragments. Consequently, while it affects approximately 14% of individuals of European descent, the impact is particularly notable among the ∼0.5% of individuals who are homozygous for the variant allele. Specifically in this group we noted increased odds of inconclusive results and decreased accuracy of the commonly used predictive models that quantify the amount of placenta-derived DNA in NIPT^14,15^. While our results only focused on the consequences for NIPT, it is very likely that homozygosity for R206C will have a large effect on liquid biopsies in oncology and other applications as well^16-18^.

Genotyping this variant in samples as part of these assays would be a first step towards sample stratification. However, many clinical applications of cfDNA make use of untargeted low-coverage whole-genome sequencing data, which prevents genotyping this SNP directly from the generated data itself. Although haplotype-based genotype imputation from low-coverage whole-genome sequencing performs reasonably well to make inferences about the R206C genotype^19,20^, the lower plasma cfDNA concentration, and thus limited cfDNA sequencing yield, complicates imputation. Our previous work showed that haplotype-based imputation of the R206C genotype resulted in a genotyping error of approximately 10 percent^13^. Furthermore, beyond the R206C genotype, other genetic^5,21^ and non-genetic^22-24^ sources of altered DNase1L3 have been reported, including immune-system related factors and chemicals such as heparin, which may have similar consequences for cfDNA.

Here we show that supervised machine learning algorithms can use the fragmentomics properties of cfDNA to accurately identify R206C homozygote samples. This method makes much more accurate predictions than genotype imputation, works with as few as 10.000 sequenced fragments, and is applicable to both paired- and unpaired sequencing data.

The computational approaches presented here can be applied to unseen samples and are an important step towards identifying plasma samples with impaired DNase1L3 activity. This will be of benefit for the ever-increasing number of cfDNA-based assays, independent of the aim of these assays.

## Methods

### Study Population

We analyzed low-pass whole-genome sequencing data from 129,676 plasma samples generated for routine non-invasive prenatal testing (NIPT) in the Amsterdam UMC as part of the Dutch TRIDENT-2 study^25^. Written informed consent was obtained from all participating women. Approval for the study was granted by the Dutch Ministry of Health, Welfare, and Sport (license 1017420-153371-PG) and the Medical Ethical Committee of VU University Medical Center Amsterdam (No. 2017.165). Samples in which aneuploidies were detected, and samples from women that did not consent to the use of their data for scientific research were excluded from the analyses. For an in-depth characterization of the studied population of pregnant women, we refer to van der Meij et al. 2019^25^.

The first training set consisted of left-over plasma from a genotype stratified collection of 63 non-invasive prenatal tests. Digital droplet PCR was used to obtain the maternal genotype, as previously described in Linthorst et al^13^. For our second dataset, an additional 106 samples with discordant genotypes between our predictor and haplotype imputation, were validated using the same protocol. As a result, both datasets were enriched for R206C homozygote plasma samples and did not represent a random sampling of the population.

### Genotype Imputation using haplotypes

Haplotype based genotypes of NIPT samples were imputed using QUILT version 0.1.9 and the Human Reference Genome Consortium dataset^26^. For details we refer to Linthorst et al 2024^13^.

### Fragmentomics feature extraction

Insert-size distributions, cleave-site motifs and 5’ end patterns were extracted from NIPT samples using a command line utility called cfstats (https://github.com/jasperlinthorst/fragmentomics), which uses pysam (https://github.com/pysam-developers/pysam) for parsing SAM files. Cleave-site motifs were defined as the bases on the GRCh38 reference assembly in a 4bp window around the 5′ extremes of a paired-end aligned cfDNA fragment. For computational efficiency, we stored only the lexicographically smallest value between the forward and the reverse-complemented sequence of each motif. The 4bp at the 5’ ends of cfDNA fragments were used as 5’ ends features. Fragment-sizes were derived from the insert-size feature in pysam. Counts were normalized to frequencies for each analysis. We used the following filters for samtools in processing our aligned sequencing data with -F3852 and -q60.

### Machine learning and dimensionality reduction

Various supervised machine learning classifiers were trained and validated using the scikit-learn package.^27^ The unsupervised UMAP^28^ implementation (https://github.com/lmcinnes/umap) was used to reduce fragmentomics features to a two-dimensional space, subsequent visualizations were generated using matplotlib^29^. We used UMAP with 100 neighbors and 500 epochs to obtain two-dimensional representations from the input data. Reference data and the prediction models are available through a web api and can be accessed through the cfstats package (https://github.com/jasperlinthorst/fragmentomics) and an online demo application (https://huggingface.co/spaces/jasperlinthorst/cfstats-demo) to apply our models on external data (either NIPT or other cfDNA liquid biopsy sequencing data).

### Classifier training and allele frequency inference

An initial Linear Discriminant Analysis (LDA) classifier was trained using 1392 cfDNA features (from 63 ddPCR genotyped samples. Feature vectors contained the following frequencies; fragment sizes in the range from 0 to 1000, 256 4bp 5’ end patterns and 136 4bp cleave-site motifs. This first cohort consisted of 22, 21 and 20 samples with wildtype, heterozygote and homozygote alleles. An additional cohort was obtained by randomly selecting 106 samples in which LDA and haplotype imputation disagreed on the R206C allele. Genotypes in this cohort were obtained with the same ddPCR protocol (see Linthorst et al 2024^13^). Random Forest (RF), Neural Network (Multi-layer Perceptron: MLP) and Support Vector Machine (SVC) classifiers were trained on the aggregate of both datasets, consisting of 169 samples. Genotypes were binarized by labeling homozygotes for the variant as 1 and homozygous wildtype and heterozygotes as 0. R206C allele frequencies were derived from classifier predictions by taking the square root of the frequency of predicted homozygotes in the entire population of 129,676 women.

## Results

### Supervised fragmentome predictors enable recognition of R206C homozygotes

We previously demonstrated a strong association between the R206C genotype, imputed from low-coverage whole-genome sequencing (lcWGS) data, and various alterations to the fragmentome^13^. However, our validation experiments indicated that ∼10% of the imputed genotypes were incorrect. All incorrect predictions were related to misclassifications between heterozygosity and homozygosity for the R206C variant. Given the much larger effect-size estimates in our validation dataset, than in our imputation dataset, we argued that it might be feasible to infer R206C genotypes from the fragmentomic properties alone, circumventing the need for genotype imputation.

To evaluate this, we used our existing dataset of 63 ddPCR-genotyped samples (22 homozygous variant, 21 homozygous wild-type, 20 heterozygous) to train a binary linear discriminant analysis (LDA) classifier using three main global fragmentomic properties as input: fragment sizes, cleave-site motifs and 5’ end patterns.

Applying the model to our dataset of 128,672 NIPT samples that were available at that time, we observed high concordance between the LDA-predicted and imputed homozygosity: 128,111 (99.6%) samples matched, and 561 (∼0.4%) were discordant. Of these, the LDA predicted 383 (∼2/3) to be homozygous variant but were imputed as either homozygous wild-type or heterozygous (type 1 discordant), and 178 (∼1/3) showed the inverse pattern (type 2 discordant). Targeted genotyping by ddPCR of 106 randomly selected discordant samples (76 type 1, 30 type 2) confirmed the fragmentomics-based prediction in 90% (95/106, all of which were type-1 errors, where the classifier predicted samples to be homozygous for R206C, but in practice were not) of the cases, indicating improved classification of R206C homozygotes using a simple LDA on fragmentomic features over haplotype imputation in this setting.

By adding the additional 106 labeled samples we obtained a dataset of 169 labeled plasma samples enriched for boundary cases, containing 21 homozygous wt, 64 heterozygous and 84 homozygous variant plasma samples. This final labeled dataset was used to retrain additional classifiers (LDA, random forest (RF), support vector classifier (SVC), neural network: multilayer perceptron (MLP)). Performance of the different classifiers was evaluated using five-fold cross-validation. Although no large performance differences were observed, the random forest model, with an average accuracy of 0.97 and AUC of 0.98 across the folds, achieved the highest accuracy (Figure 1A).

**Figure 1.**
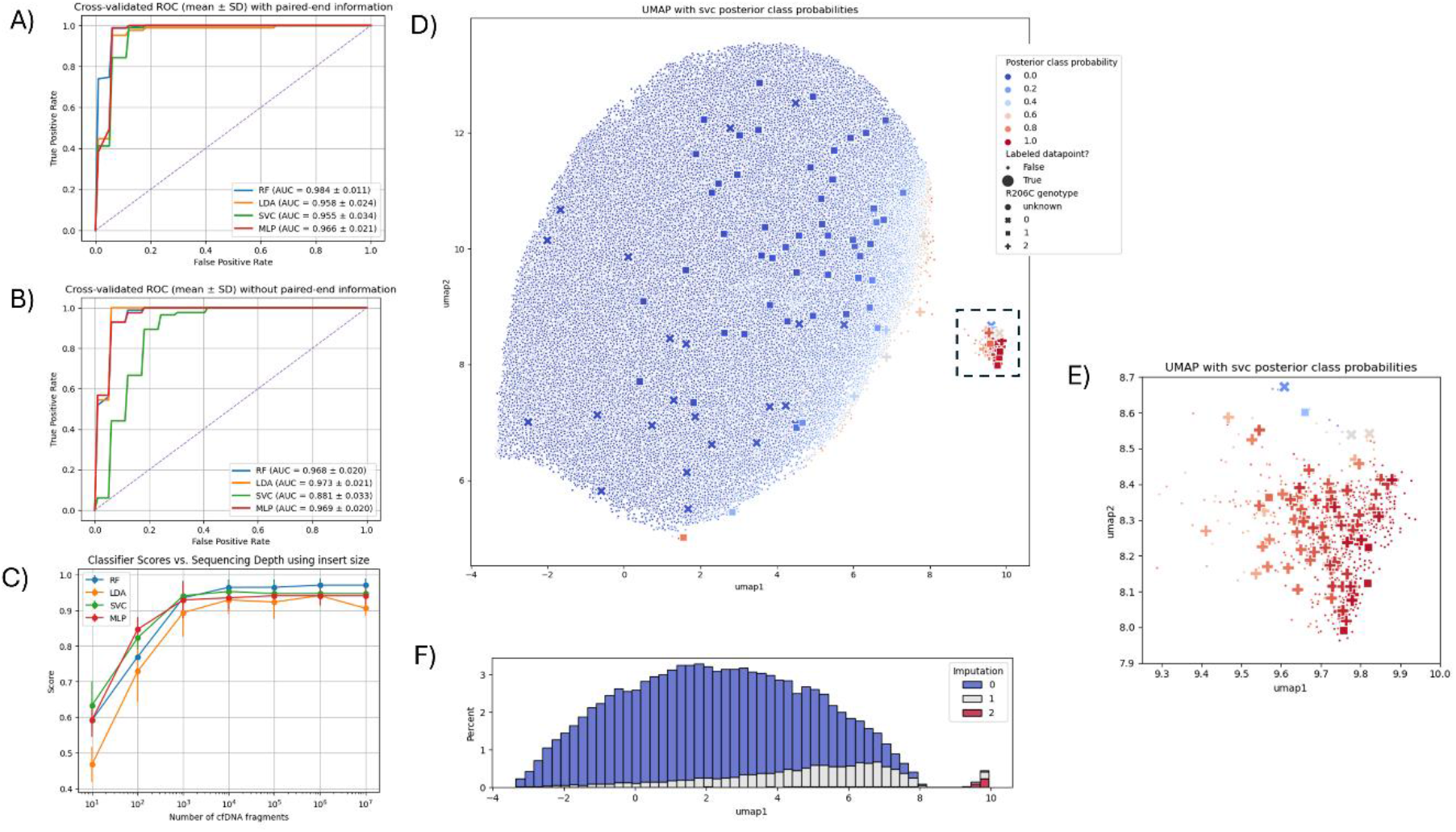
A) Mean cross-validated ROC curves for classification based on paired-end sequencing data. Curves represent the mean across 5 stratified folds. B) same as A, but without using the paired-end information, so only based on cleave-site and 5’end motifs C) classifier performance measured by AUC (y-axis) across different sequencing depths (x-axis). errorbars indicate standard deviation across 5 folds. D) Visualization of a 2 dimensional UMAP representation of the 1032 fragmentomic features across 129,676 samples, enables near-perfect separation of R206C homozygotes. Rectangle highlights the R206C homozygote cluster. ddPCR genotyped samples have large points, the shape of these points corresponds to ddPCR genotype. The color corresponds to the posterior class label probability of the SVC. E) Zoomed in visualization of the R206C homozygote cluster selected in D. F) Stacked histogram of haplotype imputed genotypes across the first UMAP dimension.

### Accurate classification can be achieved with minimal sequencing data without the need for paired-end information

Many predictors make use of fragment size which requires paired-end or long-read data. As these data increase costs and are not always available, we investigated whether cleave-site and 5’end motifs alone could also predict the R206C genotype. Using the distribution of frequencies of 4-bp sequence motifs, a random forest classifier was able to distinguish homozygote samples with only a slightly lower average AUC of 0.97 (Figure 1B).

To assess the minimum sequencing depth required, we down-sampled the validation dataset and evaluated classifier performance. Using paired-end inferred fragment sizes, classification retained high accuracy with as few as 1,000 fragments per sample (Figure 1C), while models that only used unpaired features required ∼100-fold more reads to achieve comparable performance. This is still well within the typical sequencing depths of several million fragments needed for NIPT and other types of cfDNA analysis.

### All classifier predictions deviate from population allele frequency estimates for R206C

Despite high cross-validated performance, the number of predicted homozygotes varied widely when different classifiers were applied to the full cohort. We attributed this to the relatively small size and composition of the training set, in which the homozygous allele was overrepresented with respect to the entire population. We previously estimated the R206C allele frequency in our population using the aggregate of all our data to be approximately 7%. Using Hardy-Weinberg equilibrium we therefore estimated that our dataset of 129,676 samples should approximately contain ∼846 homozygote samples. Conversely, we were able to derive allele frequency (AF) estimates from the predictions of the trained classifiers. While all classifiers predicted more R206C homozygote samples than expected, we found that, with a derived AF of ∼9.8%, the support vector classifier (SVC) most closely approximated the expected AF for R206C.

### Visualization of decision boundaries in lower dimensional space reveals a tight cluster of R206C homozygotes

We used UMAP to project the fragmentomes of all samples onto a two-dimensional space. In this space, a strongly enriched cluster of R206C homozygotes (the R206C cluster) could clearly be identified (Figure 1D). By labeling samples with the posterior class probability obtained from the trained classifiers, we were able to visualize approximate decision boundaries within this UMAP space. This visualization revealed that, although all classifiers performed comparably on training data, their predictions on the unlabeled cohort varied substantially. The SVC produced decision boundaries that best aligned with the observed homozygote cluster and most closely reproduced population R206C allele frequencies. However, by manually classifying samples on the basis of their 2d coordinates in UMAP space (e.g. all samples with the first UMAP dimension larger than 8.5), we obtained an even closer approximation of the R206C population allele frequency with an estimate of 7.7%.

To investigate how the different fragmentomic features contributed to the lower dimensional space, we also applied UMAP on the three different feature types separately (Figure S1). Although insert size features were most informative for the identification of the R206C cluster, only upon combining all three feature types a perfectly separable cluster emerged (Figure 1D/E).

### Unsupervised clustering confirms discordant cluster assignment of misclassifications

Despite our close approximation of the expected allele frequency by manually classifying samples in this lower dimensional representation (Figure 1E), investigation of the obtained clustering revealed that this approach would result in a total of 14 (8.3%) misclassified samples (9 type1, 5 type 2), significantly more than our trained supervised classifiers, which employ more complex, but potentially overfitted, decision boundaries. We observed 9 samples that clustered with R206C homozygotes but lacked the homozygous genotype (Figure 1E). Of these, 5 had the heterozygous and 4 the homozygous wildtype genotype. Conversely, 5 ddPCR confirmed homozygotes were observed close to, but outside the typical R206C cluster. We found that these samples had intermediate cfDNA size distributions, with a limited relative increase and decrease in multi and mono-nucleosome size fragments respectively, when compared to the population of WT and heterozygotes. Due to the large overlap in fragmentation profiles between R206C heterozygote and wildtype individuals, and therefore limited clinical use of differentiating between them, we did not pursue classification between these two classes. However, when the frequency of the error-prone imputed genotypes was plotted along the first UMAP dimension, we observed a highly significant effect for the heterozygous allele as well (Figure 1F).

### Longitudinal data indicate an indirect effect of R206C on the fragmentome and a role for non-genetic factors

Our plasma samples were collected over a four year period in which many participating women had more than one pregnancy for which they had NIPT. Through unique identifiers we traced the 755 plasma samples assigned to the R206C cluster to 682 unique women. For 61 (∼9%) of these women we were able to obtain NIPT sequencing data for more than one pregnancy. Although the large majority of these samples were consistently classified by all models as R206C homozygotes in subsequent pregnancies, we observed that, when projected on the 2d UMAP space, 10 samples visually crossed the cluster assignment boundary (UMAP dimension 1 > 8.5) in subsequent pregnancies (see Figure 2).

**Figure 2.**
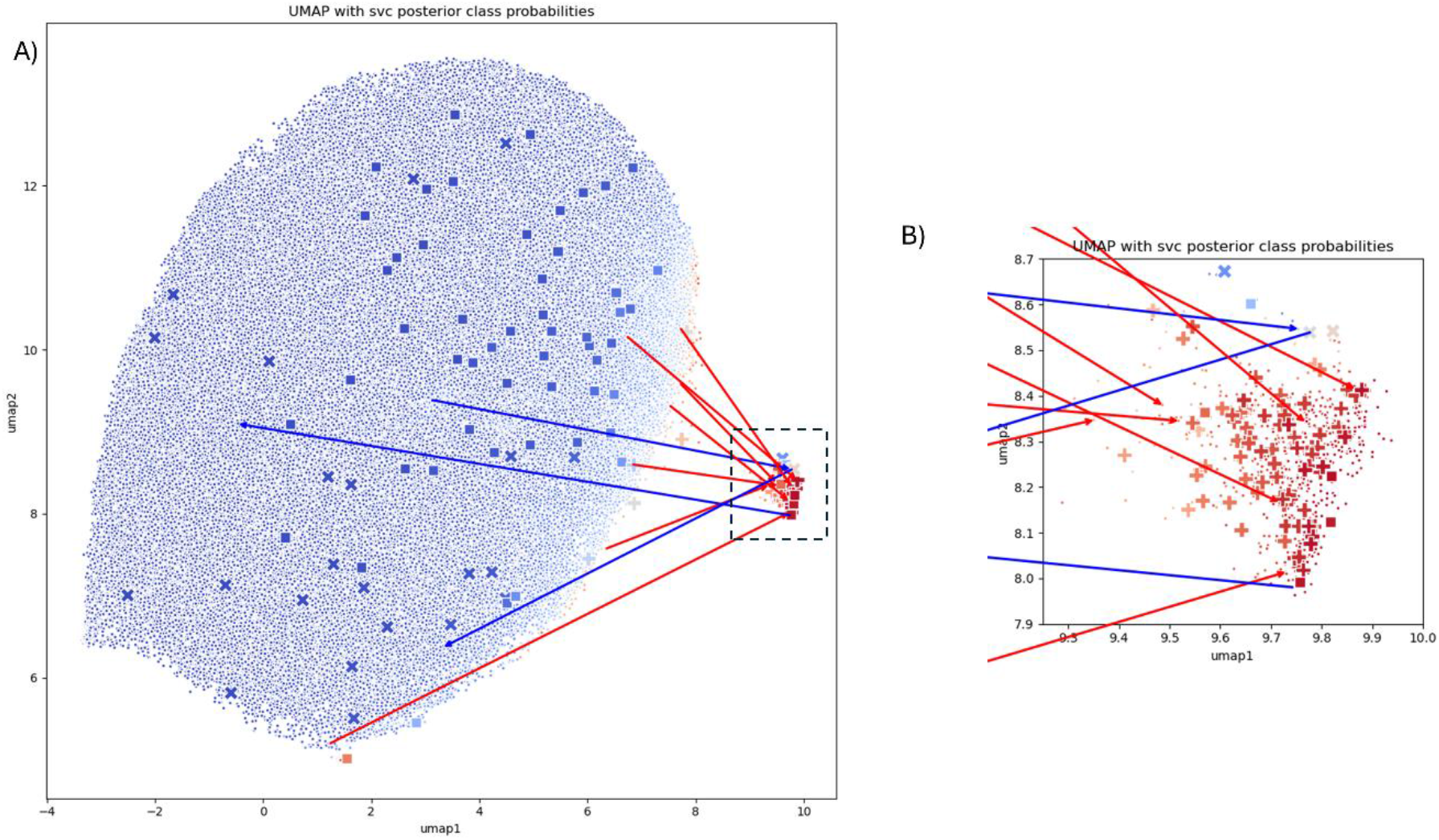
A) Same UMAP visualization as in Figure 1D, but arrows indicate the chronological progression of samples with discordant cluster assignments between subsequent pregnancies. Arrow color indicates the ddPCR validated genotypes (red=homozygous, blue=WT or heterozygous). All 7 homozygous samples move towards the R206C cluster, while heterozygous and WT samples move in both directions. B) Zoomed in visualization of the R206C homozygote cluster selected in A.

For 6 women one biobanked plasma sample was available, for four women we had biobanked samples from both pregnancies. We performed additional ddPCR analyses for these plasma samples to validate the maternal R206C genotype. Analyses resulted in identical genotypes across the four paired samples (two homozygous R206C and two wildtype), minimizing the potential for sample swaps as a possible explanation. Seven women were found to be homozygous for R206C, while three were homozygous wildtype. None were heterozygous. For all homozygous R206C samples, it was the first plasma sample in time that was not assigned to the R206C cluster, while this assignment changed to the R206C cluster in the next pregnancy. This suggests that the alterations to the ‘fragmentome’ in R206C homozygotes may accumulate over time. For the three women with the wildtype allele, we did not observe this constraint. Two were assigned to the R206C cluster only on the first test, and one only on the second test. We did not find other significant differences in potential covariates, such as age or bmi between women in or outside the R206C cluster.

## Discussion

We previously demonstrated that in a Dutch population of pregnant women the R206C variant in DNase1L3 is a major genetic determinant of variation in cell-free DNA (cfDNA). This was based on imputed genotypes from ultra-low coverage sequencing. While genotype imputation remains a useful tool, the limited accuracy—especially among homozygous individuals with lower cfDNA yields—motivates alternative approaches.

Because of the potential clinical consequences of homozygosity for this variant, we set out to develop an approach to recognize these samples from the aberrant fragmentome itself. Our data indicate that with a fraction of the sequencing input, supervised and unsupervised models trained on fragmentomic features outperform haplotype-based imputation when it comes to identifying R206C homozygotes. These insights provide a first step towards identifying R206C homozygote samples in clinical cfDNA applications.

Interestingly, our efforts indicate that despite the strong, well-recognizable effect of R206C, we are unable to obtain perfect separation of homozygous individuals. The existence of discordant cases—wildtype or heterozygous individuals displaying an aberrant fragmentome, and homozygotes lacking it—suggests that the R206C genotype alone is insufficient to fully explain the observed alterations to the fragmentome. Longitudinal analysis reinforces these observations, as in R206C homozygotes, the aberrant fragmentome appears to develop over time. This suggests that either DNase1L3 activity in these individuals may deteriorate gradually or episodically, or we observe a downstream effect of the impaired DNase1L3, which accumulates over time. These findings are reminiscent of studies in mice, where conditional DNase1L3 knockouts in macrophages do not induce anti-DNA antibodies in young mice immediately, but they do develop them later in life^30^.

We speculate that, although the most obvious biological explanation would be that impairment of DNase1L3 is the direct cause of the altered fragmentome, our results suggest that it may (also) be a consequence of long-term exposure to impaired clearance. Because the release of microparticle-associated DNA is the primary target of DNase1L3, we pose that in R206C homozygotes, DNA accumulates in microparticles, which are incompletely recovered from plasma, causing a distorted view of the fragmentome, low cfDNA plasma concentrations, limited sequencing yield and repeated NIPT failures. Future work should further test the validity of these hypotheses.

In contrast, our observation that similar fragmentome alterations and their constitutions over time are observed in individuals without the genotype, suggests that the accumulation of microparticle DNA and/or its incomplete recovery from plasma, may also be a consequence of an epigenetic or environmental trigger, which either modulates the function of DNase1L3 directly or mimics the downstream effect of it. Alternatively, altered preanalytical conditions could also play a role, but this is unlikely as all samples were handled strictly according to the same protocol and in a largely automated manner.

These observations suggest that, beyond R206C genotype stratification for diagnostic purposes, the developed classification schemes in practice identify a latent biological state that is linked to the impairment of DNase1L3. Although it is currently unclear whether this state underlies pathology, immune dysregulation, or exposure to environmental stressors, our work shows that the models developed here can accurately identify it by a relatively small amount of cfDNA sequencing data. While complete loss-of-function of DNase1L3 has been linked to a specific rare form of SLE, it was recently found that up to 1/3 of all sporadic SLE cases have autoantibodies against DNase1L3 (ref Gomez-Banuelos 2023 affinity mutaration). This suggests that our models might also be of use in the early detection of systemic autoimmune diseases, such as SLE, Rheumatoid Arthritis and Systemic Sclerosis.

Our work has several limitations. Models were exclusively built on data from pregnant women generated by the Illumina Veriseq approach. Although the biological phenomenon we observe will also be detected by other platforms, transfer methods^31^ that account for the inherent technical biases of different cfDNA platforms will be needed. The same holds for use in non-pregnant and male individuals. Another limitation of our work is that we did not address heterozygosity for R206C, which affects ∼14% of individuals in our population. Although the impact of the heterozygous allele on the individual fragmentome is much smaller, from a population screening perspective, its impact may overall be much larger.

In summary, we demonstrate that cfDNA fragmentomic features can be leveraged to infer DNase1L3 activity directly from sequencing data, enabling robust identification of individuals with impaired nuclease function. Beyond the potential to improved cfDNA-based diagnostics, this approach reveals a latent biological state that may reflect broader dysregulation of extracellular DNA clearance. Future work integrating clinical phenotypes and longitudinal sampling will be essential to determine whether this fragmentomic signature can serve as an early biomarker of immune dysfunction.

## Notes

https://github.com/jasperlinthorst/fragmentomics

### Competing Interest Statement

The authors have declared no competing interest.

### Summary of Updates

Minor changes. Inclusion of Figure S1 and extension of Fig 1. Aded reference to demo and github repository

