## Supplementary Figure S1 for "A machine learning approach to infer DNase1L3 activity from plasma cell-free DNA fragmentomics"

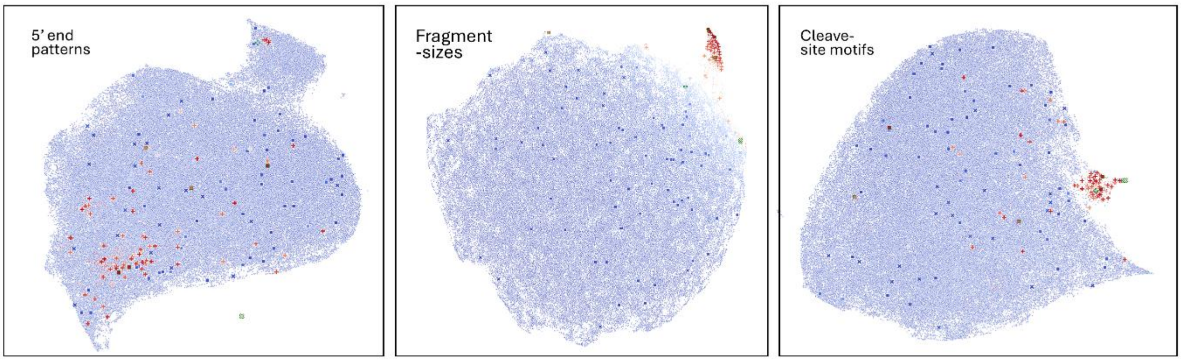


Visualization of a 2 dimensional UMAP representation of the three different fragmentomic feature types across 129,676 samples. ddPCR genotyped samples have large points, the shape of these points corresponds to ddPCR genotype. The color corresponds to the posterior class label probability of the SVC classifier trained on all features combined.
